# Tuning the brakes – Modulatory role of transcranial random noise stimulation on inhibition

**DOI:** 10.1101/2023.11.09.565862

**Authors:** Alekhya Mandali, Flavie Torrecillos, Christoph Wiest, Alek Pogosyan, Shenghong He, Diogo Coutinho Soriano, Huiling Tan, Charlotte Stagg, Hayriye Cagnan

**Affiliations:** MRC Brain Network Dynamics Unit, Nuffield Department of Clinical Neurosciences, University of Oxford, Oxford, United Kingdom, OX3 9DU; Department of Psychology, University of Sheffield, Sheffield; Neuroscience Institute, University of Sheffield; Department of Bioengineering, Imperial College London

**Keywords:** Cognitive control, transcranial random noise stimulation, beta activity, intermittent bursts

## Abstract

**Background:** Everyday decision-making requires the ability to flexibly modify and sometimes terminate our actions, such as avoiding a tempting slice of cake to hitting the brakes in an emergency. Neural oscillations, such as beta-band rhythms observed over the medial prefrontal cortex(mPFC), help regulate these context-dependent behaviours. However, how random noise stimulation would modulate neural rhythms and corresponding inhibitory behaviors remain understudied.

**Objectives:** To target the mPFC using random noise stimulation and modulate neural activity underlying inhibitory behaviours.

**Methods:** Using a single-blinded within-subject design, fifteen participants received random noise or sham stimulation in a pseudo-randomized order while performing a Go/Conflict/No-Go task. We measured neural activity and behavior before and after stimulation.

**Results:** We show that random noise stimulation significantly improved inhibitory behaviors (4.6±4.42percent) by reducing the number of errors in No-Go trials. This improvement was a function of participants’ impulsivity-levels and baseline performance, i.e., impulsive individuals who made more baseline errors improved more after receiving stimulation. At the neural level, we show that random noise stimulation increases low-beta power at stimulation site, mediated by an increase in the duration of intermittent beta-bursts.

**Conclusion:** We show for the first time that random noise stimulation improves the ability to withhold response to unexpected inhibitory cues as a function of an individual’s impulsivity level. This improvement could be attributed to increased low-beta band power and intermittent-burst duration. These results suggest that random noise stimulation of the mPFC could potentially be used as a neuromodulatory intervention to target maladaptive behaviors in impulse control disorders.

## Introduction

Cognitive control is an executive function that governs our ability to learn, modify and update actions flexibly[1, 2]. This function however could be compromised in neuropsychiatric disorders such as Parkinson’s disease and depression[3-5], and remains difficult to restore with invasive and non-invasive brain stimulation approaches. Though still under debate, it is argued that inhibitory control falls under the broad umbrella of cognitive control[6]. The medial prefrontal cortex(mPFC) is one of the critical structures implicated in this executive function[2, 7, 8], with neural activity captured as evoked response potentials(ERP) and rhythms correlating with various aspects of behavior[7-9]. Brain stimulation could be used to target specific neural dynamics, untangling the link between behavior and neural activity patterns, and may potentially yield novel approaches that could be leveraged in the future to restore cognitive control.

Neural oscillations in the mPFC capture cue-specific behavior at theta[7, 10], beta[7] and high-gamma bands[11, 12]. A recent study by Xiao et al., compared the variations across neural signatures while performing different cognitive control paradigms (e.g., Stroop and Flanker), and showed that evoked neural oscillations can be task-specific[12]. Zavala et al. reported an increase in the theta and beta bands during inhibitory trials(No-Go) and highlighted an increase in theta coherence between the subthalamic nucleus and mPFC for trials invoking cognitive control(Conflict and No-Go trials). Critically, this study suggested that the increase in the beta band power in the mPFC captured the difficulty level in a given trial, with the highest increase being observed for No-Go trials, when a complete inhibition of movement was required[7]. Beyond neural rhythms, Yamanaka et al., showed an increase in inter-trial phase-locking related to negative-positive (N2/P3) ERP components over the F_Z_ during No-Go trials[8]. Additionally, the P3 has been shown to reflect response inhibition[13] in terms of peak and onset latency as measured by stop signal reaction time tasks[14].

Transcranial random noise stimulation(TRNS) uses a wide range of frequencies(0.1-640Hz) to modulate neural activity and behavior. Previous studies demonstrated its impact on various functions: for instance, TRNS has been shown to affect working memory[15], visual perception[16, 17] and emotion perception, with age dependent effects[18, 19]. Moreover, stimulation of specific brain regions has been shown to be crucial for achieving desired behavioral effects: stimulation of the dorsolateral prefrontal cortex(dlPFC) during cognitive control improved participants’ response time[20] and could induce sustained effects following repetitive stimulation sessions[21]. On the other hand, stimulation of the inferior frontal gyrus showed no behavioral effects[22], highlighting the importance of the precise stimulation site.

Understanding the mechanism of action of stimulation is essential for optimizing TRNS applications. A study by Chaieb et al. indicated that TRNS increased cortico-excitability, an effect potentially mediated by gamma-aminobutyric acid(GABA_A_) as evidenced by the suppression of TRNS-induced cortico-excitability following GABA_A_ agonist uptake while glutamatergic receptor manipulations did not yield any changes[23]. However, how these changes in cortico-excitability reflect in neural activity and behavior across different nodes contributing to cognitive control has remained unclear. A recent computational work by West et al. suggested that changes in excitability levels mediated by lamina-specific interneurons could manifest as an increase in beta power and intermittent beta burst duration[24].

Despite the involvement of the mPFC in cognitive control, its response to TRNS during inhibitory control has been understudied. Building on previous experimental and computational work[23-27], we hypothesized that TRNS targeting the mPFC would selectively modulate inhibitory control through GABAergic mechanisms reflected as a change in the beta power and intermittent burst characteristics. To test this hypothesis, we delivered TRNS while recording participants’ neural activity using electroencephalogram(EEG) from a cohort of healthy participants as they performed a modified version of the Go/No-Go task that incorporated a conflict component (Figure 1)[7].

**Figure 1.**
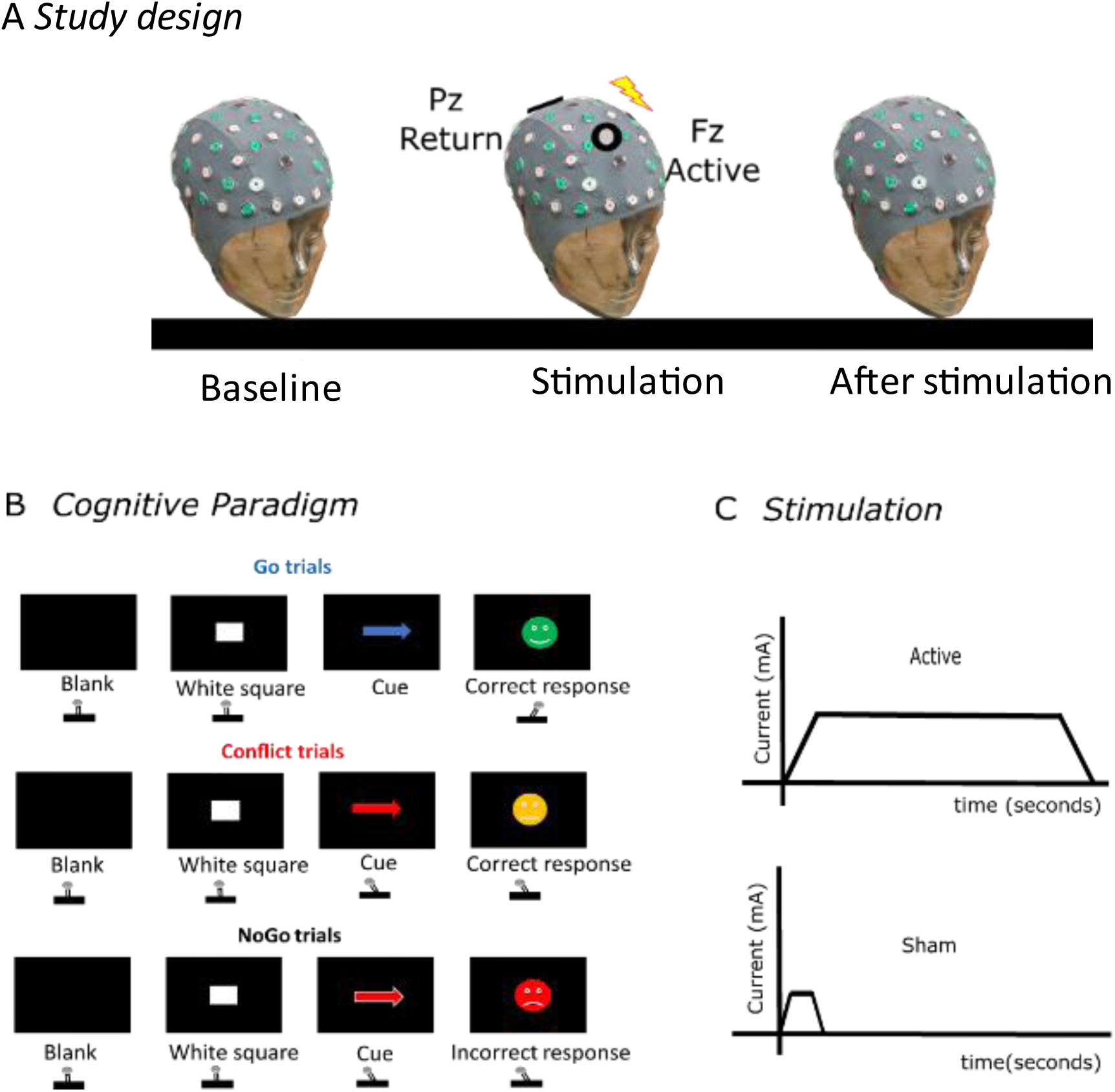
shows a summary of the experimental set up and the cognitive paradigm. (A) shows the sequence of steps in a given session measuring EEGs and behavior at baseline, during and after-stimulation (roughly within 5 minutes of completing the stimulation). TRNS was delivered with the active electrode over F_z_ and the return at P_z_. (B) shows the sequence of events during the cognitive paradigm for Go, Conflict and No-Go trials with feedback for correct, slow-correct and incorrect responses respectively and (C) indicates the stimulation pattern including the ramp-up and ramp-down of current for active (up to 1200 seconds) and sham stimulation conditions (20 seconds).

## 2 Methods and Materials

### 2.1 Participant recruitment

We recruited 16 participants from the general population through posters, and mailing lists. Participants were screened before attending the first appointment for any contra-indications for stimulation, including but not limited to a personal or family history of seizures, epilepsy, implanted electronic devices and metal in the upper body. One participant dropped out of the study due to time constraints. All participants had normal or corrected vision and were right-handed. The Central University Research Ethics Committee of the University of Oxford approved the study(CUREC-R77362/RE003). The study was undertaken in accordance with the Declaration of Helsinki, and informed written consent was obtained from all participants.

### 2.2 Experiment design

The study followed a within-subject single blinded design, during which participants received either active or sham TRNS. In each session, the sequence of events, as outlined in Figure 1A, were followed. Participants were prepped for the placement of stimulation and recording electrodes. First, participants completed a short version of the cognitive task(up to 6 minutes; 2.3 Cognitive Paradigm). TRNS was then delivered while the participants completed the longer version of the cognitive paradigm (up to 20 minutes). The participants either received active(approximately 20 minutes with ramp up/down time of 20 seconds) or sham(turned off after 20 seconds with ramp up/down time of 8 seconds) stimulation. The effect of stimulation was tested using the short version of the task(up to 6 minutes) within 5 minutes of delivering stimulation(TRNS or sham). Active and sham recording sessions were separated by at least 4 days to ensure that there were no residual stimulation effects. Participants completed questionnaires on demographics, impulsivity(UPPS measure)[28] and stimulation perception[29]. The stimulation perception questionnaire was completed after each recording session.

### 2.3 Cognitive Paradigm

We utilized a modified version of a Go/No-go task(Figure 1B) which included a conflict component[7]. Each participant was seated comfortably in a chair and the task was presented on a 28-inch computer monitor(Dell-Corporation) positioned 50cm at eye-level. Each trial started with the presentation of a blank screen for a variable duration of 1.75 – 2seconds. This was followed by a white square (0.5seconds) which indirectly served as an indicator that subsequently a cue would be presented. The cue could represent Go(arrow in blue), Conflict(arrow in red) or No-Go(arrow in blue or red with white outline), presented in a pseudo-randomized order. Response to the presented cue was measured using a joystick held using their right hand. Participants were expected to respond by moving the joystick in the direction(horizontal-movement) of the arrow for Go, the opposite direction for Conflict and not to respond for No-Go trials. Feedback was provided after a response was made, i.e., a green smiley for correct, red for incorrect and yellow for a slow response (>1second) (Figure 1B). This feedback was displayed for 0.5 seconds. Arrow direction was counter-balanced across the 3 cues controlling for direction effects. The task was coded using Psychtoolbox in MATLAB(version MATLAB 2018b, Mathworks, USA) and run on a Dell-Precision-5550(16GB, Intel i7) Windows-10 system.

The proportion of cues was 3:1:1 for Go, Conflict and No-Go, enabling the paradigm to catch uncommon events(Conflict and No-Go). The shorter version of the task had 100 trials(baseline and after stimulation) and the longer version had 300 trials(during stimulation). All participants were instructed to respond as quickly as possible at the beginning of a given session.

### 2.4 Modulation using stimulation

TRNS was delivered using a battery powered stimulator(DC-Stimulator-PLUS, NeuroConn GmbH, Germany) via conductive rubber electrodes positioned over F_Z_(Active-ring-4.8cm outer and 2.4cm inner-diameter) and P_Z_ (Return-Rectangular:5×7 cm^2^) respectively(Figure 1A). The ring electrodes allowed for measurement of neural activity at F_Z_. Each participant’s F_Z_ and P_Z_ positions were estimated using the standard 10-20 system. Before placing the stimulation electrodes, the scalp was scrubbed and prepared to improve conductivity. We applied the Ten20-conductive paste(Weaver and Company, USA) on the stimulation electrodes prior to placing them on the designated positions. Impedance was checked before placing the EEG cap(TMS-International, Netherlands) while paying particular attention to prevent bridging between the stimulation electrodes and EEG electrodes. The stimulation amplitude was titrated based on each participant’s feedback on tolerance and comfort whilst keeping the amplitude within the allowed safe limits.

### 2.5 EEG recording

EEG recordings were acquired using an EEG cap and a 32 channel TMSi-Porti amplifier (TMS-International, Netherlands). EEG recordings were limited to 16-channels spanning over the frontal, fronto-central, dominant motor cortex, and mastoid electrodes as they cover the brain regions expected to be engaged in this study[7]. The ground electrode was placed on the left forearm and EEG channels were common average referenced. The EEG recordings and the cognitive paradigm were synchronized using a trigger signal generated by an U3-HV-LabJack (LabJack-Corporation, USA) via the Psychtoolbox. Triggers were sent to the amplifier via Power-1401(Cambridge Electronic Design, UK). Distinctive voltage levels(0-2.5V) were used to mark different events(i.e., blank screen/white square/cue). The trigger signal, and displacement of the joystick were recorded simultaneously with monopolar EEGs using the TMSi-Porti amplifier and were all sampled at 2048 Hz.

### 2.6 Behavioral analysis

Behavioral data was analyzed offline using custom MATLAB scripts(version MATLAB 2018b, Mathworks, USA). Accuracy was determined based on actual and expected joystick movement directions. Reaction time was estimated from the joystick movement: data was epoched such that each segment started 1.5 seconds before and ended 1 second after cue-onset. Reaction time was then estimated as the time point when the joystick position crossed a threshold. The threshold was set at the summation of the mean baseline noise before the cue-onset and 20% of the normalized peak[30]. Baseline noise was included to account for trial-by-trial fluctuations. Trials with response times less than 100ms (i.e., premature, or accidental responses) were excluded from further analysis. Due to high intra-individual variability, all joystick movements were visually inspected and trials with premature movements and direction reversals were excluded. The reaction time per participant for correct and incorrect trials was calculated by averaging across corresponding trials. The participants with no errors were excluded from the error-trial analysis.

### 2.7 EEG Preprocessing and Analysis

#### 2.7.1 Preprocessing

EEG data was pre-processed offline using custom MATLAB(version:MATLAB-2018b, Mathworks, USA) and EEG LAB(version EEGLAB2021.1-[31]) scripts. Raw EEG data was re-referenced to the mastoid and offset corrected for drifts in the signal. The data was first band pass filtered(high-pass at 0.1 Hz and then low-pass at 100 Hz). Line noise in the signal was removed using an open-source plugin-Zapline(version zaplineplus1.1-[32]). Filtered EEG data was subjected to temporal independent component analysis[33] as implemented in EEG LAB[31]. Components representing stereotypical artifacts such as eye blinks and saccades were identified by manual inspection in both time and frequency domains and were removed from the data. On average 2.3±0.2 independent components were removed per participant. Remaining components were then back projected to obtain artifact free channel data. Data was labelled as correct and incorrect per cue type(Go/Conflict/No-Go). Finally, data was visually inspected and segments with muscle artifacts were identified and excluded from further analysis.

Labelled and cleaned data was chunked into epochs using ERPLAB(version:erplab8.3.0[34] according to the cue type (Go,Conflict and No-Go) and corrected with respect to the average 1 second [-1.5 to -0.5 before cue-onset] inter-trial interval (ITI-blankscreen). These corrected epochs will be referred to as ‘*ITI-corrected epochs*’ in the subsequent sections. The total length of each epoch was 4 seconds which included 2 seconds before the cue-onset and 2 seconds after. This epoch length was chosen to minimize boundary effects during spectral analysis and was long enough to estimate lower frequency components with sufficient time resolution.

#### 2.7.2 Time-Frequency analysis

The ‘*ITI-corrected epochs’* were then processed to calculate spectral power using Fieldtrip in MATLAB[35]. To ensure there were no boundary effects, the spectral power was calculated using a Hanning taper where 6 cycles were chosen per time window for frequencies ranging from 4 to 80 Hz at a resolution of 2 Hz for the time period of -1 to 1.2 seconds (where time=0 corresponds to cue onset) using the 4 second ITI-corrected epochs. The average power across trials was calculated per participant to identify event related changes (i.e., either event-related synchronization or event-related desynchronization compared to ITI).

#### 2.7.3 Burst analysis

We studied intermittent properties of the task-evoked oscillations in the beta band using previously validated approaches[36, 37]. First, the peak beta frequency was identified after removing the aperiodic (1/f) component. An opensource package (FOOOF, version:1.0[38]) was used to identify and remove the aperiodic (1/f) component from the power spectrum. Participants with no distinct oscillatory activity in the beta band were excluded from further analysis.

Individual peak frequency was then used to bandpass filter the signal (±2Hz) using a second-order zero-phase lag Butterworth filter. The Hilbert envelope was calculated using the continuous and cleaned EEG data which was then epoched for 4.5seconds (2seconds before and 2.5seconds after cue-onset) whilst normalizing to the ITI period. All corrected-envelope trials were concatenated to determine the burst threshold (75^th^-Percentile). It should be noted that the threshold was condition specific (here No-Go) similar to the approach used by[24]. This threshold was then used to determine the presence of bursts and the effect of TRNS on them. To identify the presence of a burst, the activity was expected to last at least one beta cycle. Estimated burst features (see supplementary:S3) were then averaged to obtain an average feature per participant.

### 2.8 Statistical Analysis

#### 2.8.1 Behavioural analysis

We ran a 2×2 repeated-measures-ANOVA for state(baseline and after-stimulation) and condition (TRNS and sham) for Go and Conflict trials for both accuracy and reaction times for correct trials. Post-hoc tests were Bonferroni corrected. For error trials, participants with no errors were excluded and mean reaction times were calculated.

Performance in No-Go trials(accuracy) was tested using a non-parametric Friedman’s 2-way analysis as the data did not pass the Kolmogorov-Smirnov normality test. Pairwise comparisons between conditions were Bonferroni corrected to adjust for multiple comparisons.

#### 2.8.2 Cluster based statistics for spectral data

Statistical difference between two time-frequency power spectrums was evaluated using a cluster-based Monte Carlo non-parametric method, using the MATLAB based fieldtrip package. Here, a paired t-test on each time-frequency combination(power-spectrum A and power-spectrum B) was compared and clusters were identified and labelled as significant if its summed t-statistic value exceeded 97.5% of the randomized distribution(p < 0.025) to test for both positive and negative differences. Neighboring channels for each of the 15 EEG channels were calculated using a triangulation method (See[35] for details) resulting in an average of 4.9 neighbors per channel. The procedure was repeated 3000 times while randomly exchanging labels between the 2 power-spectrums for each subject between for the 0-1second time period.

##### 2.8.2.1 Comparing non-cue and cue-evoked activity at baseline

The power spectrums corresponding to the non-cue period (1second before cue-onset) and cue period (1second after cue-onset) were first computed and extracted using Fieldtrip. We then collapsed the power spectrum corresponding to Go, Conflict and No-Go events similar to the approach used in [7] and statistically compared them (non-cue vs cue periods) using the cluster based approach.

##### 2.8.2.2 Effect of stimulation on Cue-specific activity

The effect of stimulation on spectral power was estimated by comparing the power spectrums corresponding to baseline condition and after-stimulation between 0 and 1seconds after cue onset and frequency range 8-20Hz. This frequency range was selected based on the cluster range observed while comparing baseline and cue-evoked power spectrums(Fig 2C,2D).

**Figure 2.**
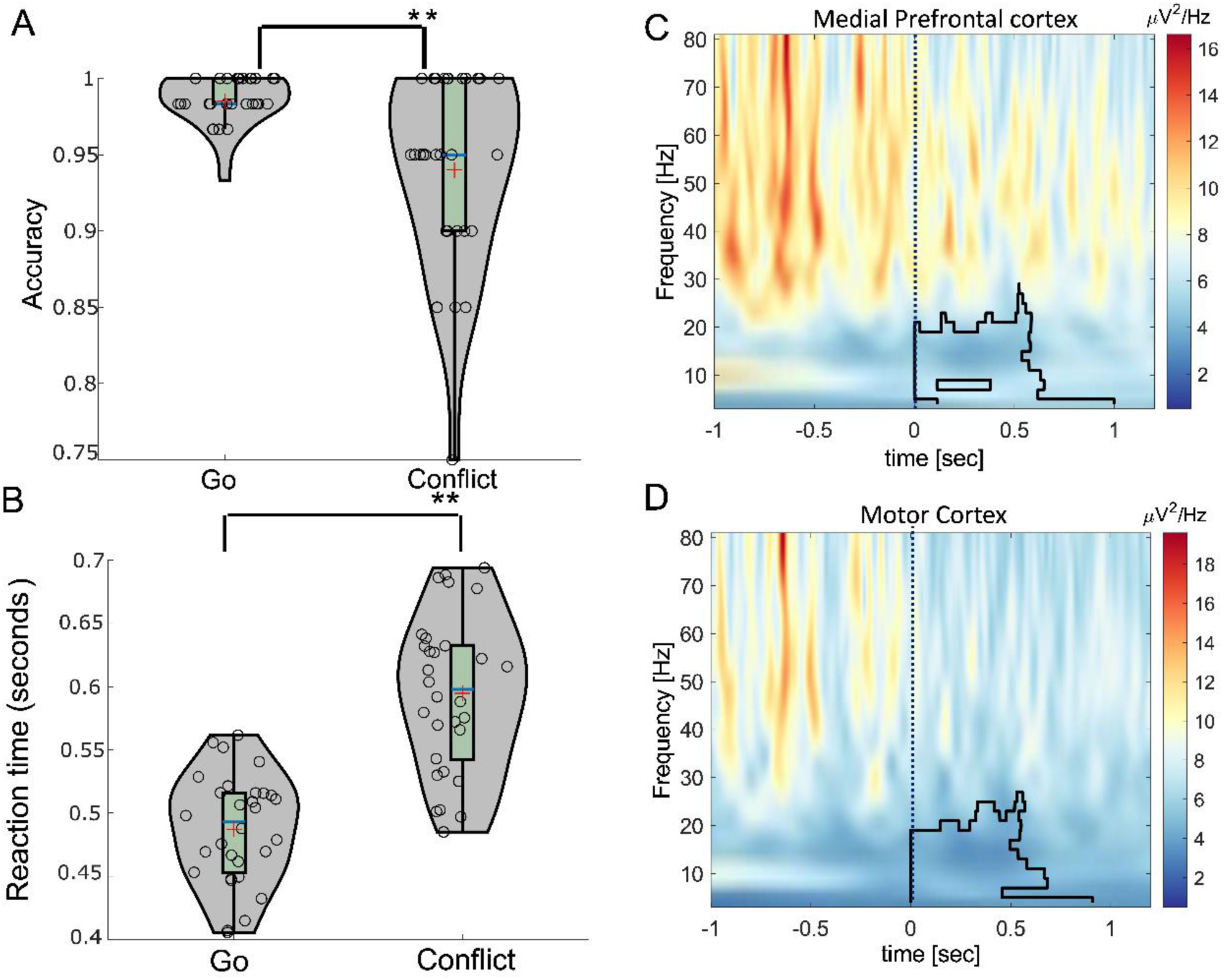
shows the baseline behaviour and evoked activity. (A) and (B) indicate the baseline accuracy and reaction time for correct Go and Conflict trials collapsed across baseline TRNS and sham sessions and (C) and (D) show the spectral powers evoked due to presentation of a cue when collapsed across Go, Conflict and No-Go trials for baseline TRNS session. The outline corresponds to significant clusters computed using the Montecarlo non-parametric test (p< 0.025) when comparing evoked activity between [0 1] seconds with non-cue period of the same window length [-1.0 0] seconds where ‘0’ is cue-onset. The red cross in (A) and (B) indicates the mean of the sample and blue horizontal bar indicates the median. ** indicates p<0.001.

#### 2.8.3 Burst Analysis

Various burst features such as duration, amplitude, and number of bursts were compared by calculating the median across trials for each participant for F_Z_ and C_3_. Extracted values were then compared using a 2×2 ANOVA(baseline and after stimulation) for TRNS and sham conditions. Pairwise comparisons between conditions were Bonferroni corrected to account for multiple comparisons.

## 3 Results

### 3.1 Demographics

Participants (6-female, 9-male) were non-smoking, aged 25.8±6.04years and had an impulsivity score of 38.5±7.8[28]. The mean current(mA) was 1.58±0.44 for TRNS and 0.76±0.3 for sham sessions, respectively. The average impedance was under 10kΩ (9±5.15kΩ) across both sessions. The stimulation perception questionnaire[29] collected after each session indicated the presence of expected sensations such as itching and fatigue at moderate levels. The Bang-blinding-index[39] for active (BI=0.2) and sham (BI=-0.13) sessions indicated an acceptable level of blinding.

### 3.2 Baseline spectral activity and behaviour

We first validated our cognitive paradigm’s ability to evoke expected neural activity and behaviour(accuracy and reaction time) in line with the previous literature[7].

#### 3.2.1 Baseline behavior

We first evaluated behavior at baseline (before receiving stimulation) by calculating the accuracy and reaction time(seconds) during Go and Conflict trials.

We compared accuracy and reaction times during Go and Conflict trials at baseline by collapsing across TRNS and sham conditions. As expected, accuracy for the conflict condition(0.94±0.06) was significantly lower (t(29)=3.72, *p*<0.001) than Go(0.98±0.01) (Figure 2A). Similar to accuracy, we observed a significant difference between Go and Conflict reaction times(t(29)=-16.29, *p*< 0.001): trials involving conflict(0.59±0.06) were significantly slower than Go(0.49±0.04) (Figure 2B).

#### 3.2.2 Baseline vs Evoked Time-frequency Analysis

We compared the spectral powers evoked due to presentation of a cue when collapsed across Go, Conflict and No-Go trials for baseline TRNS session compared to non-cue period. In line with the previous literature, we observed an increase(See Supplementary Figure-S1) in spectral power in theta and decrease in lower-beta bands(p<0.005) over F_Z_(Figure 2C) and C_3_(Figure 2D) after the onset of a cue(Go/Conflict/No-Go), compared to the non-cue period.

### 3.3 Effect of TRNS on neural activity and behaviour

#### 3.3.1 Effect of stimulation on Go/Conflict behaviour

We next explored the effect of stimulation during Go and Conflict trials on accuracy and reaction times (2×2 repeated-measures-ANOVA– state: baseline and after stimulation and condition:TRNS and sham).

For accuracy, as hypothesized, there was no main effect of state (F(1,14)=0.49, p=0.49) or condition (F(1,14)=1.39, p=0.26) or an interaction (F(1,14)=0.02, p=0.88) in Go trials (Figure 3A). For Conflict trials (Figure 3B), there was a main effect of state (F(1,14)=13.82, p=0.002) however no main effect of condition (F(1,14)=0.5, p=0.7) or interaction (F(1,14)=0.02, p=0.89). A Bonferroni corrected post-hoc t-test for state showed an increase in accuracy after stimulation (irrespective of active or sham) compared to baseline (mean ± std error: baseline= 0.94±0.01, after-stimulation=0.98±0.01, p=0.006). This indicates the effect of practice on Conflict trials.

**Figure 3.**
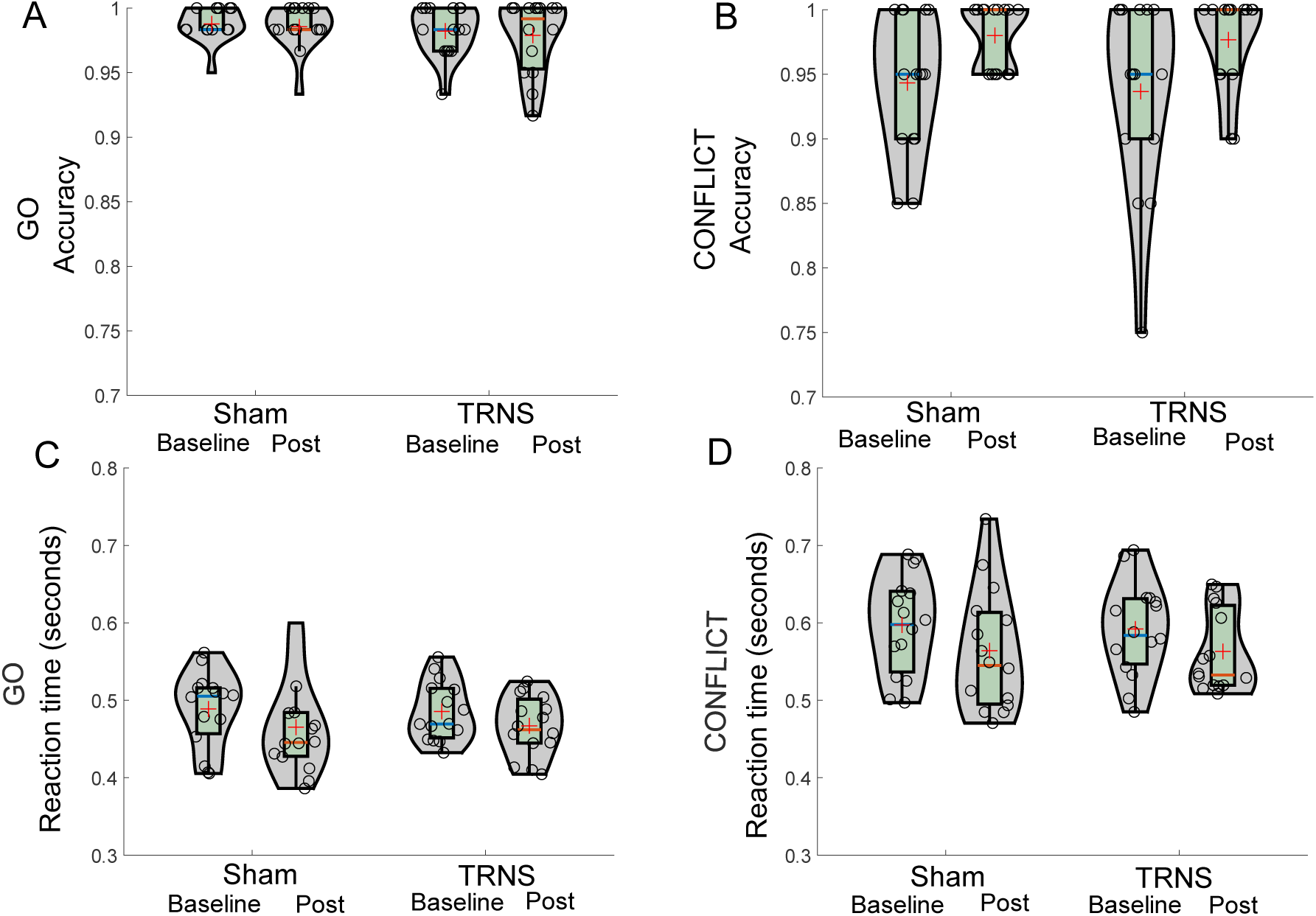
shows the accuracy and reaction time (seconds) during baseline and after-stimulation (post) for sham and transcranial random noise stimulation (TRNS) for correct Go (A &C) and correct Conflict (B & D) trials. The red crosses and blue horizontal bars on the violin plots indicate mean and median, respectively.

Considering the differences in stimulation amplitude across participants, we used the average current(sham and TRNS) as a covariate to the 2×2 ANOVA. We only found an interaction effect between condition and current amplitude (F(1,13)=6.17, p=0.027) and no other significant effects during Go and Conflict trials. Despite the interaction effect there was no correlation between participants’ accuracies and the current amplitude.

Similarly, for reaction time, there was a main effect of state(F(1,14)=7.55, p=0.02) but no main effect of condition(F(1,14)=0.02, p = 0.92) or an interaction(F(1,14)=0.09, p=0.76) during correct Go trials(Figure 3C). A Bonferroni corrected post hoc t-test for state showed a significant decrease in reaction time (p=0.02) after stimulation (irrespective of active or sham)(0.47±0.01) compared to baseline(0.49±0.01). For correct Conflict trials(Figure 3D), there was a main effect of state(F(1,14)=17.59, p<0.001) but no main effect of condition(F(1,14)=0.08, p=0.78) or an interaction(F(1,14)=0.04, p=0.85). A Bonferroni corrected post hoc t-test for state showed a significant decrease in reaction time(p<0.001) after stimulation(active or sham)(0.56±0.02) compared to baseline(0.6±0.01). When considering the average current as a covariate, we did not observe any significant main or interaction effects for Go and Conflict trials.

These results confirmed that there was no stimulation related modulation of behaviors (accuracy and reaction time) concerning Go and Conflict trials.

#### 3.3.2 Effect of stimulation on No-Go related behaviour and neural activity

Since accuracies during the No-Go condition did not pass the normality test, we used a two-way Friedman’s non-parametric test to compare accuracies(Figure 4A) at baseline(TRNS-0.95±0.04, sham-0.97±0.04) and after stimulation(TRNS-1, sham-0.99±0.02) for TRNS and sham conditions. This test showed significant differences between the distributions(χ2(3)=15.8, *p*=0.001) with pairwise comparisons across conditions showing a significant increase in accuracy for only the TRNS condition(*p*=0.035).

**Figure 4.**
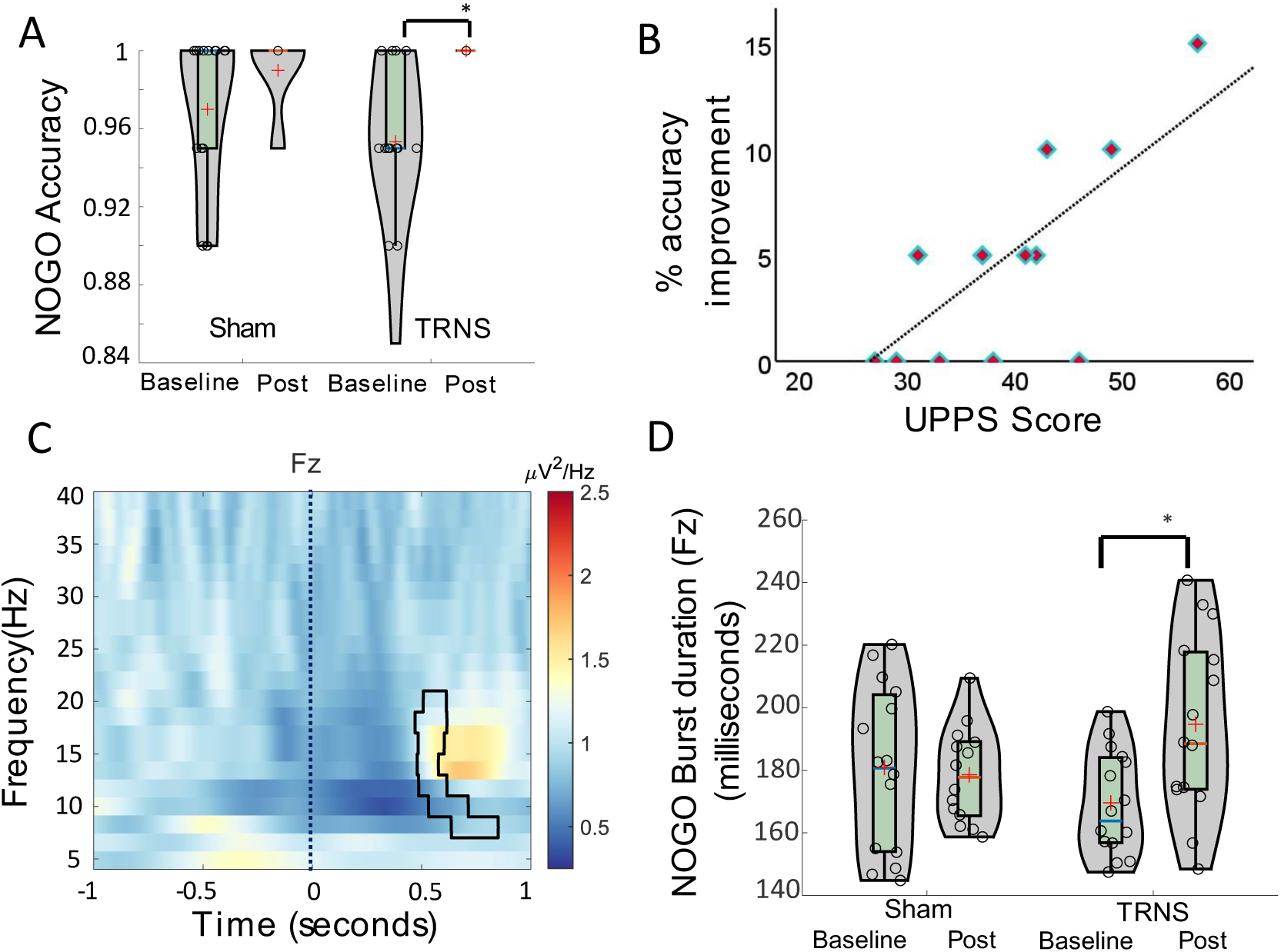
shows the accuracy and spectral powers for No-Go trials. (A) shows the accuracy levels at baseline and after-stimulation for Sham and TRNS conditions. (B) shows the improvement in the individual accuracies as a function of their impulsivity scores after TRNS and (C) shows the F_Z_ spectral power after TRNS. The outline shows the increased power in the time-frequency domain when comparing baseline with after TRNS and (D) shows the average burst duration changes across F_z_. The outline in Plot C indicates the significant cluster (p<0.025) and the dotted line indicates the onset of the Nogo cue. * indicates p<0.05.

We also studied the relationship between participant specific impulsivity levels and inhibitory behaviours(No-Go accuracy). A non-parametric Spearman’s correlation showed an inverse relationship between the baseline No-Go accuracy and impulsivity UPPS scores(ρ=-0.51, *p=0*.02), i.e., individuals with higher impulsivity scores made more errors in baseline-TRNS condition. Furthermore, a Spearman’s correlation showed a positive correlation(Figure 4B) between the impulsivity scores and percentage improvement after TRNS (ρ=0.57, *p=*0.03), i.e., individuals with higher impulsivity scores had better improvement in their accuracy scores after TRNS but not after sham (ρ=-0.43, *p =* 0.1). We further studied if the improvement in accuracy was correlated with the stimulation amplitude. A non-parametric Spearman’s correlation showed no relationship for TRNS(ρ=0.0653, *p=*0.81) or sham(ρ=-0.25, *p=*0.37) conditions.

To further explore neural signatures driving this improvement in No-Go accuracy, we compared the spectral power over the F_z_ and C_3_ corresponding to the No-Go trials (baseline and after stimulation for TRNS and sham conditions) after cue-onset. TRNS stimulation increased the spectral power in the beta band (*p=0.022)* over F_z_(cluster highlighted with an outline in Figure-4C) between 0.5 and 1 seconds after cue-onset(time=0) compared to baseline. This increase in spectral power was absent in the sham condition. We didn’t observe any significant changes in beta power over C_3_ for TRNS or sham conditions.

##### 3.3.2.1 Changes in Average Burst features in No-Go condition

To further understand the change in spectral power(Figure 4C) after TRNS, we extracted intermittent beta-burst features (average amplitude, average duration, and total number of bursts) at baseline and after stimulation for both TRNS and sham conditions. One participant was excluded from this analysis due to lack of a distinct beta peak. The features corresponded to average metrics per participant(Figure:S2,S3).

###### Fz- Burst features

For burst duration (milliseconds, Figure 4D), there was a main effect of state(F(1,13)=6.36, p=0.025) but not condition(F(1,13)=0.16, p=0.69) or interaction(F(1,13)=8.91, p=0.011). A Bonferroni corrected post hoc t-test for state showed a significant increase in burst duration(p=0.025) after stimulation(TRNS or sham)(187.35±4.82) compared to baseline(174.81±4.36). A paired sample t-test also highlighted a significant increase in burst duration after TRNS(t(13)=-4.5, *p*<0.001) but not sham (t(13)=0.32, *p*=0.75). For burst amplitude, there was no main effect of state (F(1,13)=3.46, p=0.085) or condition(F(1,13)=0.125, p=0.73) or interaction(F(1,13)=0.115, p=0.74). Similarly, for the total number of bursts, there was no effect of state(F(1,13)=0.2, p=0.66) or condition(F(1,13)= 0.15, p=0.7) or interaction(F(1,13)=1.17, p=0.3).

###### C_3_- Burst features

For burst amplitude, there was no main effect of state(F(1,13)=0.82, p=0. 38) or condition(F(1,13)=0.001, p=0.98) or interaction (F(1,13)=0.03, p=0.87). Similarly, for burst duration, there was no main effect of state (F(1,13)=1.17, p=0.29) or condition (F(1,13)=0.06,p=0.8) or interaction(F(1,13)=0.01 p=0.92) and for burst number, we observed no main effect of state(F(1,13)=2.73, p=0.12) or condition(F(1,13)=0.58, p=0.45) but interaction(F(1,13)=8.27 p=0.013). A paired sample t-test showed an increase in burst number after TRNS (t(13)=-3.56, p<0.01) but not for the sham condition(t(13)=0.46, p=0.65).

The intermittent burst analysis therefore further delineates our observed increase in spectral power after TRNS stimulation (Figure- 4C) to be contributed by an increase in burst duration (Figure-4D).

## 4. Discussion

We targeted the mPFC using random noise stimulation and sham, while participants performed a cognitive control task and showed that TRNS improved inhibitory control by decreasing the error rate during No-Go events. This behavioral effect was mediated via an increase in the low-beta power over the mPFC compared to before stimulation, which was absent for the sham stimulation condition. This spectral power change at mPFC was mediated by an increase in the average burst duration of intermittent beta bursts. Critically, participants with a higher impulsivity score made more errors before stimulation and showed a greater improvement in their inhibitory control after TRNS, as reflected in their decreased error rates.

### 4.1 Effect of TRNS on inhibitory control

We show for the first time that the TRNS induced improvement in stopping behaviors (Figure 4A) as a function of participants’ baseline impulsivity levels(Figure 4B), which was absent in the sham condition. We also observed that individuals with higher self-reported impulsivity scores made more errors in withholding their response, similar to that observed in [40]. TRNS had a differential effect on inhibitory control, i.e., participants with higher impulsivity improved more(Figure 4C) after receiving stimulation. This further supports the notion that impact of stimulation on behavior could be a function of baseline performance[19, 41, 42], as observed in other stimulation techniques such as transcranial direct current stimulation [43]. These findings emphasize the importance of understanding how heterogeneity due to factors such as psychological traits, anatomy, and neurochemistry influence neuro-modulatory effects of stimulation[44].

We then show that this improvement in stopping behaviors after TRNS is mediated through an increase in the low-beta band power(low-beta up to 20Hz)(Figure 4C) over the mPFC during No-Go trials. This observed increase in spectral power coincides with the approximate reaction time during Go and Conflict trials(Figure 2C and 2D) (i.e., when a movement would be expected). We therefore argue that the observed rise in spectral power after TRNS, specifically in this time window when a movement was observed in Go and Conflict trials, may be a potential counteractive mechanism to improve inhibition during No-Go trials.

Beta was one of the two prominent bands that has been observed over the mPFC, with an ascending oscillatory power across Go, Conflict and No-Go trial [7]. Previous studies have shown that TRNS may induce changes in GABA concentration[23], which in sensorimotor cortex, has been shown to play a key role in beta rhythms in terms of peak frequency[25, 27] and amplitude[26]. While the precise mechanism through which TRNS modulates beta rhythms remains unknown, taking in to account the findings from previous work, one could argue that this modulatory effect could be driven by GABA_A_. Another potential explanation is that TRNS modulates task-related activity via repeated sub-threshold perturbations[45].

The stop signal task[46] has also been shown to measure response inhibition. Both intracranial and scalp EEG studies showed an increase in the beta activity within few hundred milliseconds after the stop signal[47]. Similarly, recordings from Parkinson’s disease patients[48] undergoing deep brain stimulation surgery of the subthalamic nucleus showed an increase in right-frontal beta, suggesting a strong association between stopping and beta activity. Schimdt et al argued that the increase in beta activity could correspond to a ‘clear out’ mechanism leading to stopping[47]. They also suggested that this mechanism was not restricted to motor activity but also could generalize to thoughts and decision making[47]. Beta rebound after a movement has also been linked to GABAergic activity and proposed as a resetting mechanism[47]. Taking in to account these findings, we argue that mPFC ‘beta’ reflects behaviors linked to inhibition.

It has recently been shown that oscillatory activity at beta and gamma bands exist as ‘bursts’, i.e., short transient cycles of activity in sensorimotor cortex[24, 36, 37, 49-52]. Here, we observed similar burst profiles over the mPFC: duration of these temporally localized intermittent bursts was increased by TRNS(Figure 4D). Previously, our research group has shown that burst features in the motor cortex could be modulated by the connectivity strength of interneurons (GABAergic): specifically, an increase in the connectivity strength of the interneurons inversely correlated with beta burst duration[24]. We therefore posit that TRNS may lead to an increase in the overall burst duration and power by modulating the complex excitatory-inhibitory connectivity of the mPFC via interneurons and GABAergic signaling.

To understand the communication between the mPFC and motor cortex which potentially contributed to the improvement in No-Go accuracy, we studied phase coherence between the mPFC and motor cortex. While we did not observe a change in phase synchrony before and after stimulation, we did observe an increase in the beta-burst number over the motor cortex which could underlie improved stopping behaviours.

### 4.2 Effect of TRNS on behaviours involving movement

Our cognitive control paradigm included Go, Conflict and No-Go trials, allowing for an evaluation of response time and accuracy during Go and Conflict trials, and accuracy during No-Go trials. As previously reported, participants had higher accuracy(Figure 2A) and were faster(Figure 2B) for trials corresponding to Go compared to Conflict trials[7]. There was no effect of TRNS on accuracy or reaction time for both Go[21] and Conflict conditions. However, there was a practice effect on the performance, particularly for Conflict trials: participants were faster and more accurate after stimulation, regardless of receiving TRNS or sham. It has been shown that the effect of TRNS strongly depended on the site of stimulation with improvements in Go reaction times when the stimulation site was dlpfc[21] versus no effect when stimulation was delivered to the right inferior frontal gyrus [22, 42, 53]. Our findings also showed no improvement in performance during Go and Conflict trials, which could be attributed to either the stimulation site (mPFC) or absence of a DC-offset during stimulation as previously used by Brever-Aeby et al.[21]. Beyond the brain area, the effect of stimulation on behaviour could be dependent on the type of paradigm, as different oscillatory bands are evoked by different cognitive tasks[12]. For example, a recent study using TRNS stimulation of the DLPFC during a Stroop Task, showed no effects on reaction time[54] unlike the improvement observed in Go/No-Go[21].

To summarise, we showed that random noise stimulation could improve inhibitory control when the mPFC is the stimulation target. In line with our hypothesis, the improvement in inhibitory control was linked to an increase in the beta power and intermittent burst duration. These results highlight the capability of TRNS to improve impulsive behaviours and could open novel avenues for treating disorders of impulsivity.

### 4.3 Limitations and Future work

Our study is not without limitations. Based on previous literature, we hypothesized that the effect of TRNS on mPFC could be GABA mediated, however this requires further research either using MRS spectroscopy or a paired pulse TMS protocol such as short interval intracortical inhibition which reflects GABAergic tone. To study how stimulation modulated the interaction between medial prefrontal cortex and motor cortex, we calculated the phase spectral index but did not find any differences. This aspect needs to be further studied. Taking into account the intra-individual variability and the effect of non-invasive stimulation, the wider effect of our protocol needs to be studied with higher number of participants and increased number trials across each of the events.

## Supporting information

Supplementary Methods and results

## Data Availability

The analysis code will be made available online.

## Acknowledgements

We would like to thank our participants for kindly taking part in this study. We would also like to thank Dr Carolina Reis and Dr Tim West for their inputs in the study. This study was funded by the Medical Research Council UK Award MR/R020418/1 and MR/X023141/1 (HC). CJS is funded by a Wellcome Trust Senior Research Fellowship (224430/Z/21/Z). SH was supported by the Guarantors of Brain (HMR04170) and the Royal Society (IES\R3\213123). HT is supported by the Medical Research Council UK [MC_UU_00003/2, MR/V00655X/1, MR/P012272/1], the National Institute for Health Research (NIHR) Oxford Biomedical Research Centre (BRC) and the Rosetrees Trust, UK.

## Author Contributions

AM: study design, data collection and analysis and manuscript preparation. FT: data collection and manuscript, CW: data collection and manuscript, AP: data collection and manuscript, SH: data collection and manuscript, DSC: data collection and manuscript, CS: manuscript, HT: manuscript, HC: study design, data analysis and manuscript

## References

[1] Braver TSJTics. The variable nature of cognitive control: a dual mechanisms framework. 2012;16(2):106-13.

[2] Miller EKJNrn. The prefontral cortex and cognitive control. 2000;1(1):59-65.

[3] Cocchi L, Harrison BJ, Pujol J, Harding IH, Fornito A, Pantelis C, et al. Functional alterations of large−scale brain networks related to cognitive control in obsessive−compulsive disorder. 2012;33(5):1089–106.

[4] Paulus MPJCOiBS. Cognitive control in depression and anxiety: out of control? 2015;1:113-20.

[5] Singh A, Richardson SP, Narayanan N, Cavanagh JFJN. Mid-frontal theta activity is diminished during cognitive control in Parkinson’s disease. 2018;117:113–22.

[6] Aron ARJTn. The neural basis of inhibition in cognitive control. 2007;13(3):214-28.

[7] Zavala B, Jang A, Trotta M, Lungu CI, Brown P, Zaghloul KAJB. Cognitive control involves theta power within trials and beta power across trials in the prefrontal-subthalamic network. 2018;141(12):3361–76.

[8] Yamanaka K, Yamamoto YJJoCN. Single-trial EEG power and phase dynamics associated with voluntary response inhibition. 2010;22(4):714–27.

[9] Nigbur R, Ivanova G, Stürmer BJCN. Theta power as a marker for cognitive interference. 2011;122(11):2185–94.

[10] Cavanagh JF, Wiecki TV, Cohen MX, Figueroa CM, Samanta J, Sherman SJ, et al. Subthalamic nucleus stimulation reverses mediofrontal influence over decision threshold. 2011;14(11):1462–7.

[11] Bartoli E, Conner CR, Kadipasaoglu CM, Yellapantula S, Rollo MJ, Carter CS, et al. Temporal dynamics of human frontal and cingulate neural activity during conflict and cognitive control. 2018;28(11):3842–56.

[12] Xiao Y, Chou C-C, Cosgrove GR, Crone NE, Stone S, Madsen JR, et al. Cross-task specificity and within-task invariance of cognitive control processes. 2023;42(1):111919.

[13] Enriquez-Geppert S, Konrad C, Pantev C, Huster RJJN. Conflict and inhibition differentially affect the N200/P300 complex in a combined go/nogo and stop-signal task. 2010;51(2):877-87.

[14] Huster RJ, Messel MS, Thunberg C, Raud LJC. The P300 as marker of inhibitory control–fact or fiction? 2020;132:334-48.

[15] Murphy O, Hoy K, Wong D, Bailey N, Fitzgerald P, Segrave RJBS. Transcranial random noise stimulation is more effective than transcranial direct current stimulation for enhancing working memory in healthy individuals: Behavioural and electrophysiological evidence. 2020;13(5):1370–80.

[16] Battaglini L, Contemori G, Penzo S, Maniglia MJNL. tRNS effects on visual contrast detection. 2020;717:134696.

[17] Van der Groen O, Tang MF, Wenderoth N, Mattingley JBJPcb. Stochastic resonance enhances the rate of evidence accumulation during combined brain stimulation and perceptual decision-making. 2018;14(7):e1006301.

[18] Penton T, Bate S, Dalrymple KA, Reed T, Kelly M, Godovich S, et al. Using high frequency transcranial random noise stimulation to modulate face memory performance in younger and older adults: Lessons learnt from mixed findings. 2018;12:863.

[19] Penton T, Dixon L, Evans LJ, Banissy MJJSr. Emotion perception improvement following high frequency transcranial random noise stimulation of the inferior frontal cortex. 2017;7(1):1–7.

[20] McIntosh JR, Mehring CJSr. Modifying response times in the Simon task with transcranial random noise stimulation. 2017;7(1):1–16.

[21] Brevet-Aeby C, Mondino M, Poulet E, Brunelin JJNC. Three repeated sessions of transcranial random noise stimulation (tRNS) leads to long-term effects on reaction time in the Go/No Go task. 2019;49(1):27-32.

[22] Brauer H, Kadish NE, Pedersen A, Siniatchkin M, Moliadze VJNp. No modulatory effects when stimulating the right inferior frontal gyrus with continuous 6 Hz tACS and tRNS on response inhibition: a behavioral study. 2018;2018.

[23] Chaieb L, Antal A, Paulus WJFin. Transcranial random noise stimulation-induced plasticity is NMDA-receptor independent but sodium-channel blocker and benzodiazepines sensitive. 2015;9:125.

[24] West TO, Duchet B, Farmer SF, Friston KJ, Cagnan HJPiN. When do bursts matter in the primary motor cortex? Investigating changes in the intermittencies of beta rhythms associated with movement states. 2023;221:102397.

[25] Baumgarten TJ, Oeltzschner G, Hoogenboom N, Wittsack H-J, Schnitzler A, Lange JJPo. Beta peak frequencies at rest correlate with endogenous GABA+/Cr concentrations in sensorimotor cortex areas. 2016;11(6):e0156829.

[26] Hall SD, Barnes GR, Furlong PL, Seri S, Hillebrand AJHbm. Neuronal network pharmacodynamics of GABAergic modulation in the human cortex determined using pharmaco− magnetoencephalography. 2010;31(4):581–94.

[27] Porjesz B, Almasy L, Edenberg HJ, Wang K, Chorlian DB, Foroud T, et al. Linkage disequilibrium between the beta frequency of the human EEG and a GABAA receptor gene locus. 2002;99(6):3729–33.

[28] Cyders MA, Littlefield AK, Coffey S, Karyadi KAJAb. Examination of a short English version of the UPPS-P Impulsive Behavior Scale. 2014;39(9):1372–6.

[29] Fertonani A, Ferrari C, Miniussi CJCN. What do you feel if I apply transcranial electric stimulation? Safety, sensations and secondary induced effects. 2015;126(11):2181–8.

[30] Szul MJ, Bompas A, Sumner P, Zhang JJBRM. The validity and consistency of continuous joystick response in perceptual decision-making. 2020;52:681–93.

[31] Delorme A, Makeig SJJonm. EEGLAB: an open source toolbox for analysis of single-trial EEG dynamics including independent component analysis. 2004;134(1):9–21.

[32] de Cheveigné AJN. ZapLine: A simple and effective method to remove power line artifacts. 2020;207:116356.

[33] Bell AJ, Sejnowski TJJNc. An information-maximization approach to blind separation and blind deconvolution. 1995;7(6):1129–59.

[34] Lopez-Calderon J, Luck SJJFihn. ERPLAB: an open-source toolbox for the analysis of event-related potentials. 2014;8:213.

[35] Oostenveld R, Fries P, Maris E, Schoffelen J-MJCi, neuroscience. FieldTrip: open source software for advanced analysis of MEG, EEG, and invasive electrophysiological data. 2011;2011.

[36] Cagnan H, Mallet N, Moll CK, Gulberti A, Holt AB, Westphal M, et al. Temporal evolution of beta bursts in the parkinsonian cortical and basal ganglia network. 2019;116(32):16095–104.

[37] Tinkhauser G, Pogosyan A, Tan H, Herz DM, Kühn AA, Brown PJB. Beta burst dynamics in Parkinson’s disease OFF and ON dopaminergic medication. 2017;140(11):2968–81.

[38] Donoghue T, Haller M, Peterson EJ, Varma P, Sebastian P, Gao R, et al. Parameterizing neural power spectra into periodic and aperiodic components. 2020;23(12):1655–65.

[39] Bang H, Ni L, Davis CEJCct. Assessment of blinding in clinical trials. 2004;25(2):143–56.

[40] Dück K, Overmeyer R, Mohr H, Endrass TJP. Are electrophysiological correlates of response inhibition linked to impulsivity and compulsivity? A machine−learning analysis of a Go/Nogo task. 2022:e14310.

[41] Van der Groen O, Wenderoth NJJoN. Transcranial random noise stimulation of visual cortex: stochastic resonance enhances central mechanisms of perception. 2016;36(19):5289–98.

[42] Varheenmaa M, Wikgren J, Thomas O, Kortteenniemi A, Brem A-K, Lehto SMJBSB, Translational,, et al. Modulation of impulsive behaviours using transcranial random noise stimulation. 2022;15(1):32–4.

[43] Fertonani A, Miniussi CJTN. Transcranial electrical stimulation: what we know and do not know about mechanisms. 2017;23(2):109–23.

[44] Li LM, Uehara K, Hanakawa TJFicn. The contribution of interindividual factors to variability of response in transcranial direct current stimulation studies. 2015;9:181.

[45] Reed T, Cohen Kadosh RJJoimd. Transcranial electrical stimulation (tES) mechanisms and its effects on cortical excitability and connectivity. 2018;41(6):1123–30.

[46] Verbruggen F, Aron AR, Band GP, Beste C, Bissett PG, Brockett AT, et al. A consensus guide to capturing the ability to inhibit actions and impulsive behaviors in the stop-signal task. 2019;8:e46323.

[47] Schmidt R, Ruiz MH, Kilavik BE, Lundqvist M, Starr PA, Aron ARJJoN. Beta oscillations in working memory, executive control of movement and thought, and sensorimotor function. 2019;39(42):8231–8.

[48] Ricciardi L, Apps M, Little SJnPsD. Uncovering the neurophysiology of mood, motivation and behavioral symptoms in Parkinson’s disease through intracranial recordings. 2023;9(1):136.

[49] van Ede F, Quinn AJ, Woolrich MW, Nobre ACJTin. Neural oscillations: sustained rhythms or transient burst-events? 2018;41(7):415–7.

[50] Feingold J, Gibson DJ, DePasquale B, Graybiel AMJPotNAoS. Bursts of beta oscillation differentiate postperformance activity in the striatum and motor cortex of monkeys performing movement tasks. 2015;112(44):13687–92.

[51] Jones SRJCoin. When brain rhythms aren’t ‘rhythmic’: implication for their mechanisms and meaning. 2016;40:72-80.

[52] Sherman MA, Lee S, Law R, Haegens S, Thorn CA, Hämäläinen MS, et al. Neural mechanisms of transient neocortical beta rhythms: Converging evidence from humans, computational modeling, monkeys, and mice. 2016;113(33):E4885–E94.

[53] Sallard E, Buch ER, Cohen LG, Quentin RJNR. No evidence of improvements in inhibitory control with tRNS. 2021;1(4):100056.

[54] Dondé C, Brevet-Aeby C, Poulet E, Mondino M, Brunelin JJNC. Potential impact of bifrontal transcranial random noise stimulation (tRNS) on the semantic Stroop effect and its resting-state EEG correlates. 2019;49(3):243–8.

