## Supplementary Methods and results for "Tuning the brakes – Modulatory role of transcranial random noise stimulation on inhibition"

**Supplementary Materials**

### Supplementary Methods

*Evoked Related potential (ERP) analysis*

The '*ITI-corrected-epochs*' were used for ERP analysis. 20 trials were randomly chosen from the Go condition to ensure comparable signal to noise ratios across all conditions (Go/Conflict/No-Go). The average ERP per participant and condition was calculated and were analyzed for variations in N2 and P3 potentials across mPFC ($F_Z$) and motor cortex ($C_3$).

*Cluster based statistics for ERP data*

Statistical difference between two time-series was evaluated using a cluster-based Monte Carlo non-parametric method, using the MATLAB based fieldtrip package. The procedure was repeated 3000 times while randomly exchanging labels between the 2 conditions for each subject between 0 and 0.5 seconds.

### Supplementary Results

*Difference between cue and non-cue periods at baseline conditions*

The following figure shows the difference in power spectral density between cue [0-1 seconds] and non-cue [-1 0] across Go/conflict and No-Go trials in baseline condition for $F_Z$ and $C_3$.

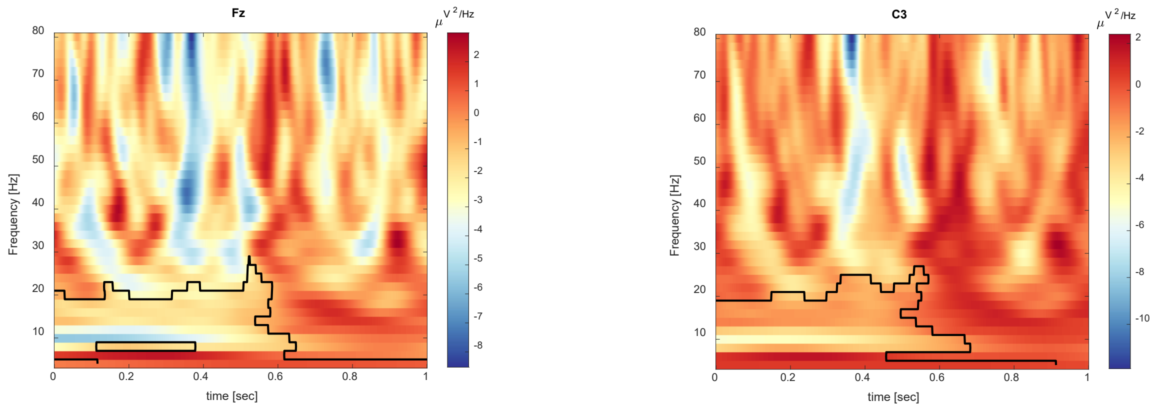


Figure S1 shows the difference in power spectral density between cue (0 to 1 second) and non-cue period (-1 0 ) in seconds.

*Effect of stimulation on behaviour*

Mean reaction times are summarized in Table S1 for Go and Conflict trials after excluding trials with premature incorrect responses for correct trials and participants with no errors for error trials. Table 1 also shows mean (± standard deviation) accuracy and reaction times (seconds) during Go and Conflict trials for TRNS and sham stimulation conditions for correct and error trials.

| Parameter | Condition | Baseline Go | After stimulation Go | Baseline Conflict | After stimulation Conflict |
|---|---|---|---|---|---|

| | | | | | |
|---|---|---|---|---|---|
| Accuracy | TRNS | 0.98±0.01 | 0.98±0.03 | 0.93±0.07 | 0.98±0.04 |
| | Sham | 0.99±0.01 | 0.99±0.02 | 0.94±0.05 | 0.98±0.02 |
| Reaction time (seconds)- Correct trials | TRNS | 0.48±0.04 | 0.47±0.04 | 0.59±0.06 | 0.56±0.05 |
| | Sham | 0.49±0.05 | 0.47±0.06 | 0.6±0.06 | 0.56±0.08 |
| Reaction time (seconds)- Error trials | TRNS | 0.52±0.14 | 0.69±0.19 | 0.64±0.12 | 0.54±0.07 |
| | Sham | 0.56±0.14 | 0.51±0.09 | 0.63±0.15 | 0.59±0.11 |

Table S1: Accuracy – minimum value is zero and maximum value is one indicating 0 and 100%, respectively.

*Effect of stimulation on burst characteristics*

The plots below show the histograms of the burst characteristics (duration) for TRNS and Sham conditions at baseline and after stimulation.

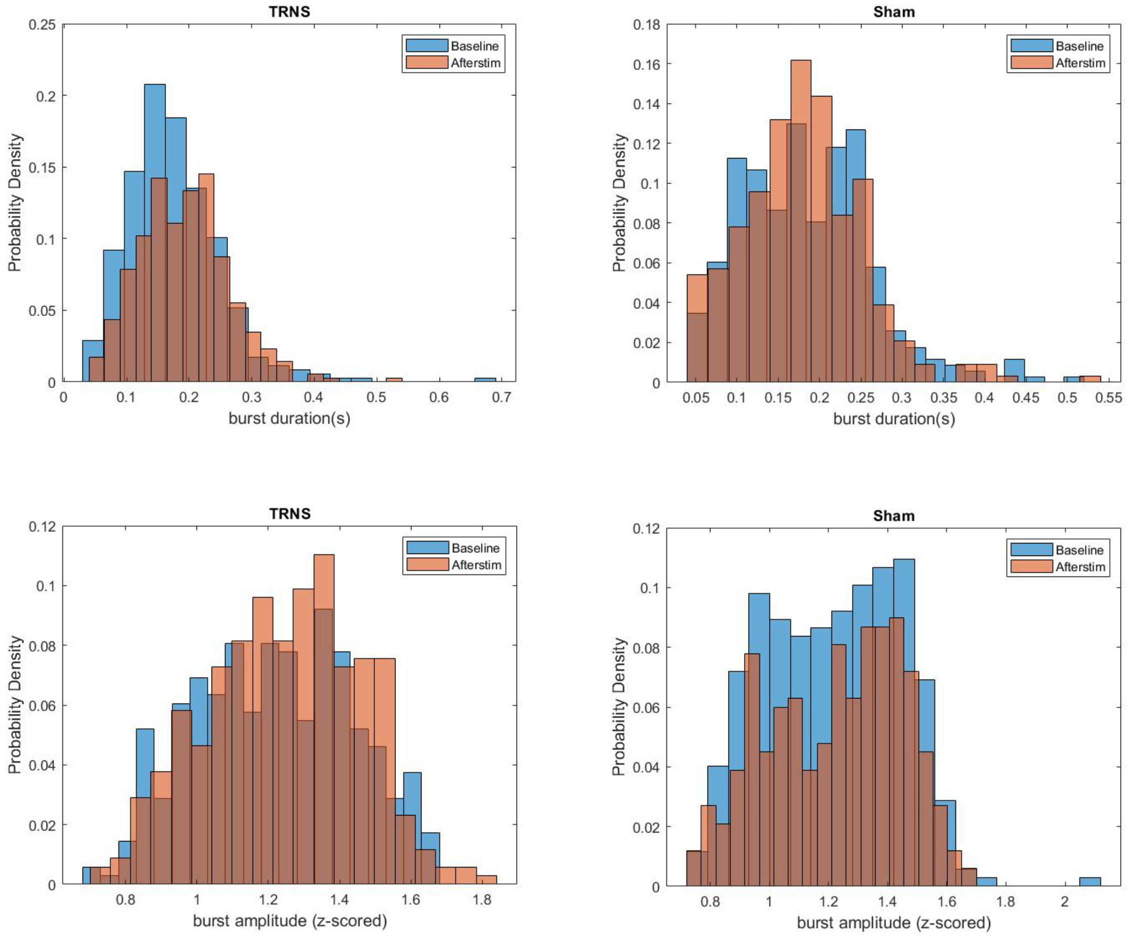

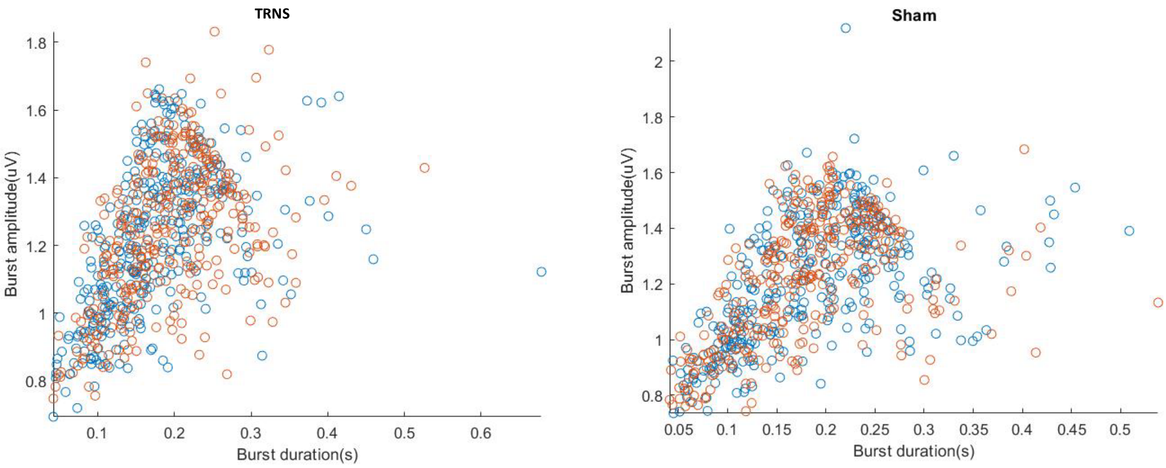


Figure S2 show the histogram of the burst duration calculated across all the participants at baseline and after stimulation for TRNS (left) and sham conditions.

Table 2 summarizes beta band burst features for $F_Z$ and $C_3$

| Parameter | Condition | Baseline $F_z$ | After stimulation $F_z$ | Baseline $C_3$ | After stimulation $C_3$ |
|---|---|---|---|---|---|
| Burst duration (milliseconds) | TRNS | 168.54±16.61 | 196.15±29.1 | 182.02±21.58 | 177±20.81 |
| | Sham | 181.1±27.3 | 178.55±15.18 | 183.37±24.1 | 179.25±19.25 |
| Burst amplitude (A.U) | TRNS | 1.22±0.06 | 1.25±0.08 | 1.24±0.04 | 1.25±0.06 |
| | Sham | 1.22±0.08 | 1.24±0.07 | 1.24±0.08 | 1.1.25±0.05 |
| Number of bursts | TRNS | 23.21±3.4 | 22.86±3.3 | 22.57±2.41 | 24.71±2.67 |
| | Sham | 23.1±2.58 | 23.86±2.65 | 23.07±2.58 | 22. 71±3.05 |
| Burst Threshold | TRNS | 0.76±0.04 | 0.74±0.03 | 0.72±0.05 | 0.74±0.05 |
| | Sham | 0.74±0.05 | 0.75±0.05 | 0.74±0.05 | 0.76±0.04 |

*Effect of stimulation on Evoked response potentials (ERP)*

'*ITI-corrected epoch'* data (see Materials and Methods) was averaged across trials per participant to obtain an average ERP which showed significant differences in the P3 potential (Figure S4). Non-parametric cluster-based statistics identified significant clusters ($p < 0.005$ after Bonferroni corrections across 6 conditions) over $F_Z$ between the Go and No-Go (Figure S4A and S4B) conditions during sham, with a higher amplitude P3 for the No-Go condition. Although we observed an effect of TRNS between No-Go and conflict (p=0.01) trials, this did not survive multiple comparisons. There was also a significant increase in P3 between Go and No-Go, and Conflict and No-Go conditions after TRNS but not sham (Figure S4C and S4D) over the motor cortex ($C_3$). It should be noted that there were no significant differences in the N2 potential.

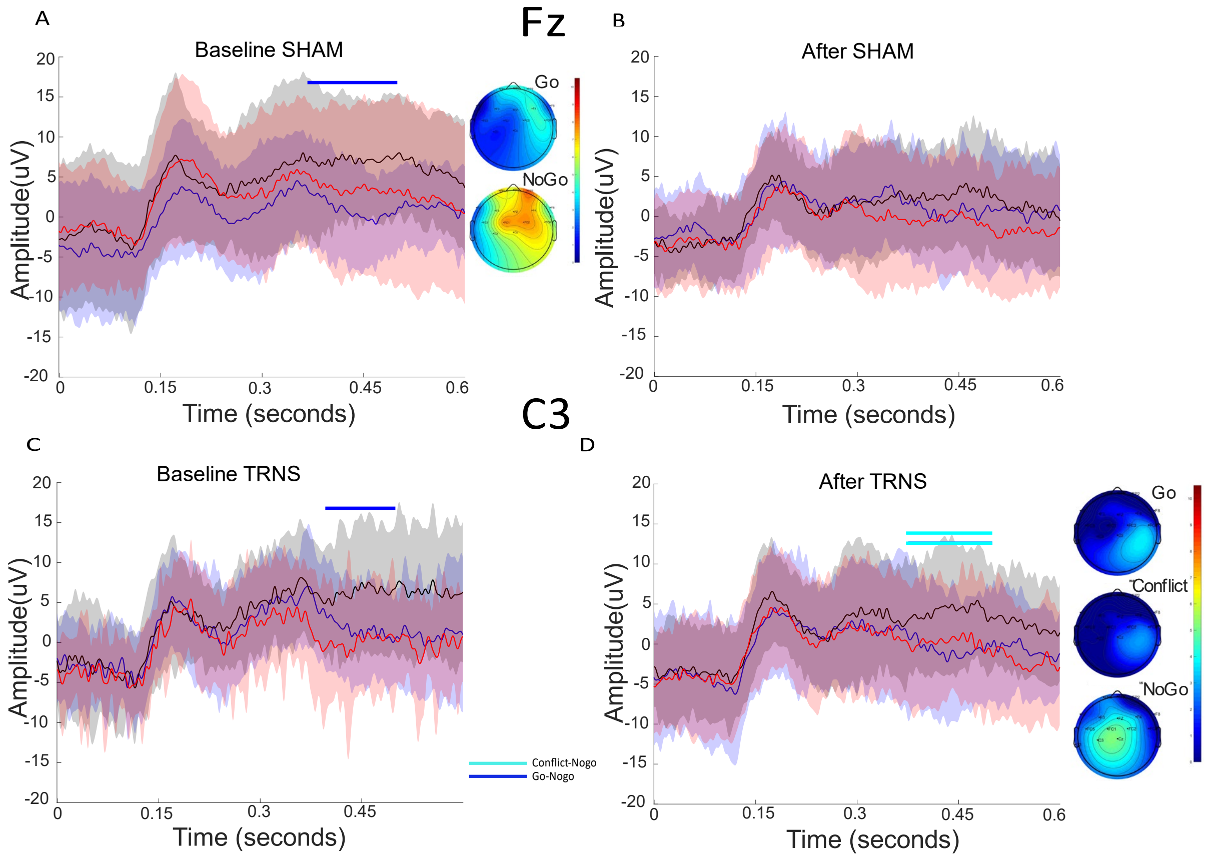


Figure S4 shows the event related potentials (ERP) across the three conditions (blue- Go, red-conflict and black-No-Go) at baseline and after-stimulation for Fz (A & B) and C3 (C & D) respectively. The x-axis corresponds to time in seconds and y-axis indicates the amplitude (μV). The bar line indicates the duration when a significant difference has been observed after correcting for multiple comparisons (Blue -Go and No-Go, Cyan-Conflict and No-Go). The topo plots in panels A & D display the average activity between 300-500milliseconds. The topo plot in panel A compares the activity between Go and No-Go condition at baseline and panel D shows Go, Conflict and No-Go (from top to bottom) after TRNS stimulation measured across C3.

For several decades, P3 potentials (occurring between 300 and 800ms after cue presentation) have been associated with rare events such as No-Go trials and known to capture inhibitory control. We observed a differential effect of TRNS on P3 across the three conditions: P3 amplitude over the motor cortex was significantly higher for No-Go compared to Go and Conflict trials after TRNS, suggesting a higher inhibitory control driving the improvement in No-Go accuracy. We did not observe any differences at the level of N2 between Go, Conflict or No-Go trials, suggesting a conflict monitoring response to all cues.
